# Deep Disentangled Representation Learning Reveals Neuron Subtype-Specific Nuclear Morphologies Across Aging in Mice and Humans

**DOI:** 10.1101/2025.10.16.682849

**Authors:** Mostofa Rafid Uddin, Zhiqian Zheng, Kashish Gandhi, Hung-Ching Chang, Jenesis Kozel, Yaswanth Gali, Silas A. Buck, Jill R. Glausier, George C. Tseng, Zachary Freyberg, Min Xu

**Affiliations:** Carnegie Mellon University, Pittsburgh, PA 15213, USA; University of Pittsburgh, Pittsburgh, PA 15260, USA; Indian Institute of Information Technology (IIIT), Dharwad, Karnataka 580009, India

**Keywords:** Imaging transcriptomics, Morphologic Profiling, Representation learning, Neural aging

## Abstract

We present a computational pipeline that links nuclear morphology to mRNA expression–based cell phenotypes under diverse biological conditions, including aging, disease progression, and drug response, using RNAscope imaging. The pipeline consists of three components: nuclear segmentation from RNAscope images, nuclear morphology identification, and downstream statistical analysis. Central to our approach is a novel unsupervised method, based on deep disentangled representation learning, which effectively captures diverse nuclear morphologies in large-scale datasets, as validated on synthetic benchmarks. We applied the full pipeline to RNAscope data targeting dopaminergic and glutamatergic neuron populations in the midbrains of mice and humans. Our analyses uncovered distinct nuclear morphology differences between dopaminergic and non-dopaminergic, as well as glutamatergic and non-glutamatergic neurons, in both species. Moreover, we identified a significant interaction between neurotransmitter identity and healthy aging in mice, reflected in systematic changes in nuclear morphology. These findings position nuclear morphology as a scalable and informative imaging-based readout of cell identity and physiological state.

## Introduction

Morphology of subcellular organelles offers important insights into their respective functions [1, 2, 3, 4, 5, 6, 7, 8]. Since the nucleus is one of the most prominent cellular organelles, its morphology has been commonly used for decades as a marker for cells for applications such as cellular quantification [9, 10, 11]. However, far less is known about the functional relevance of these nuclear morphologies. This raises a number of important questions: Do factors such as age, species, or cell type play important roles in nuclear morphology? Can distinctive nuclear morphologies be used as relevant biomarkers? Indeed, the morphology of nuclei is closely linked to processes such as cellular senescence [1], neurodegenerative diseases [12], and tumor progression [7], establishing their potential as biomarkers for biological processes and disease progression.

Answering these questions is often limited by technical challenges, including the lack of methods to effectively model morphology at a large scale (involving hundreds of thousands of nuclei). Many previous works [13] primarily relied on manual inspection of small-scale data, which potentially introduces human bias and lacks statistical confidence. Few of the prior works [1, 14] used geometric methods to estimate a small set of attributes, such as area, perimeter, moments, convexity, and aspect ratio, to quantify morphology. While these attributes provide some insights, they do not capture the holistic information on morphology. Moreover, geometric methods have low throughput and do not scale with data. Consequently, there is a need for an automated high-throughput approach.

Recent developments in AI-driven image analysis, for example, high-fidelity deep models [15, 16, 17], offer the opportunity for automated, high-throughput, unbiased approaches for morphologic quantification. This offers more comprehensive answers to the above questions, linking nuclear morphology to the aging process, region, and cell type-specific effects. The capacity for AI-driven large-scale and high-throughput analyses makes them particularly well-suited for analyzing nuclear morphology across hundreds of thousands of cells from tissue-level fluorescent images. Despite these advantages, the potential of AI-driven deep learning models in such types of research was underexplored.

In this study, we tailored a specialized deep learning method called Harmony [15] to analyze transformation-disentangled nuclear morphology across hundreds of thousands of cells across varying age groups and species, including mice and humans. After quantitative validation on a synthetic dataset that demonstrated superiority over the baseline methods, the method was applied to multiplex RNAscope fluorescence in situ hybridization (FISH) data measuring mRNA expression of key markers of dopaminergic and glutamatergic neurons in mice (n = 15) and humans (n = 12). We first segmented the nuclei from the DAPI-stained channel of the FISH images in a semi-automated manner, resulting in 1.1 million nuclei from mice and 58, 514 nuclei from humans. Using our transformation-disentangling deep learning model, we then quantified the nuclear morphology for mice and humans. Several other nuclei attributes, such as size, nucleoli count, and average DAPI intensity, were also estimated. We then evaluated differences between nuclear morphology and attributes across age groups in mice and human midbrains, as well as across cell phenotypes, including dopaminergic/nondopaminergic and glutamatergic/nonglutamatergic neurons.

Our analysis revealed significant differences in nuclear morphology between dopaminergic neurons (identified by tyrosine hydroxylase (TH) expression) and nondopaminergic neurons, as well as between glutamatergic neurons (identified by vesicular glutamate transporter 2 (VGLUT2) expression) and nonglutamatergic neurons, in both mice and humans. In the mouse midbrain, we also observed a significant interaction between neuron subtype and aging in shaping nuclear morphology. Notably, the presence of dopamine and the absence of glutamate in neurons produced similar aging-related effects on nuclear morphology. These findings offer important insights for future investigations aimed at understanding the biology of healthy aging across diverse neuron subpopulations. Additionally, our approach demonstrates the potential of linking subcellular structural morphology with image-based transcriptomics, providing a powerful tool for novel biological discovery from large-scale bioimaging datasets.

## Results

### Overview of our pipeline

In this study, we used RNAscope data with multiplex RNAscope expressions of Tyrosine Hydrolase (TH) and Vesicular Glutamate Transporter 2 (VGLUT2) from mouse and human brains throughout aging. Data were obtained from our previous work [18] with the relevant protocols followed and necessary permissions obtained. One channel of the FISH data contains 4,6-diamidino-2-phenylindole (DAPI)-stained images, which we used to determine nuclei information (Figure 1A-B). The other channels contained information from the RNAscope probe for different markers, including TH, VGLUT2, and housekeeping genes. All channels in the FISH data were aligned, thereby capturing information from the same set of cells. This enables the simultaneous determination of a cell’s phenotype (via RNAscope probes) and its nuclear properties (via DAPI stains).

**Figure 1:**
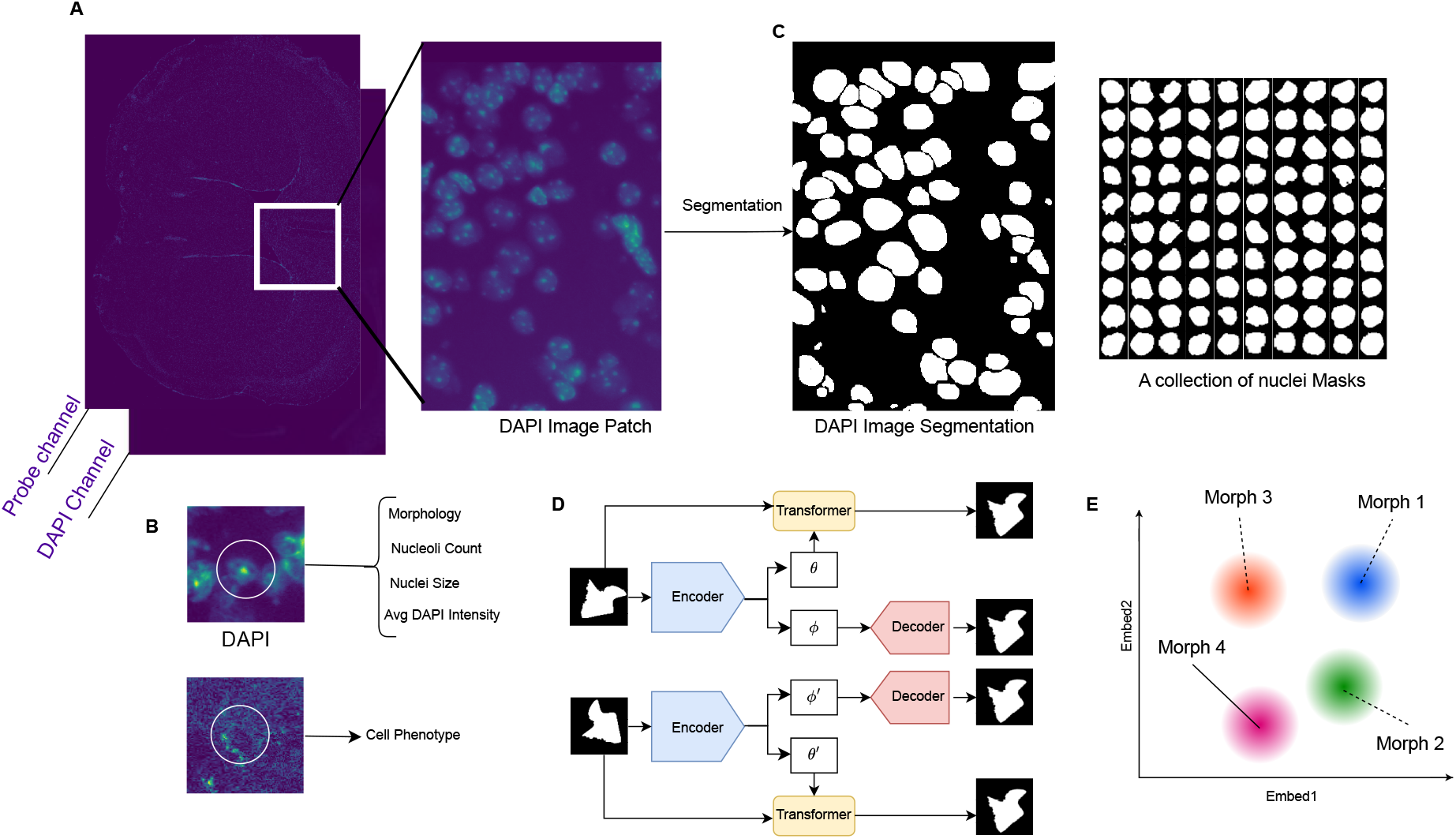
Overview of our datasets and algorithmic pipeline. **A**. The RNAscope assays in our dataset contain DAPI and probe channels. **B**. We obtain the nuclear morphology and other nuclei-associated information from the DAPI channel. From the probe channel, we obtain cell phenotype information based on grain count. To obtain nuclear information, we first segment the DAPI images patch by patch. **C**. We show our segmentation results and a collection of individual nuclei masks, which we later use for morphology identification. **D**. Our algorithm for disentangling morphology-aware semantic information from morphologyagnostic transformations. We do not show the loss for clear visualization. **E**. Morphology classification by clustering the morphology-aware semantic space.

We first segmented the nuclei from the DAPI-stained channel of the RNAscope data (Figure 1C). We observed that tools such as Halo by Indica [19] or slidebook software [20] do not properly segment many nuclei from the DAPI-stained images (Supplementary Note 2). To this end, we built our own segmentation model. We manually segmented several DAPI-stained image patches from our FISH data with LabelStudio [21]. We then fine-tuned a CellPose [11] segmentation model on these patches with our manual annotation as ground truth. We used the fine-tuned model to infer the nuclei segmentation mask for all the DAPI-stained images in our datasets, ensuring a satisfactory nuclei segmentation result.

After segmenting all the nuclei, we extracted the individual nuclei masks. We then processed the nuclei masks by removing neighboring nuclei areas, filtering erroneous segmentation, centering, and padding (details in Methods). This processing resulted in thousands of images, each visualizing the mask for a single nucleus. Each image was of size 40 *×* 40 pixels, containing a single nucleus at the image center. We then quantified the morphology of these processed nuclei masks with our specialized deep-disentangled representation learning model.

Our specialized deep learning model is based on Harmony [15], an autoencoder-like disentangled representation learning model capable of disentangling parameterized transformations from semantic contents. The morphology of an object is invariant to its orientation, shift, and scale. For example, if one nucleus can be obtained by orienting, shifting, or scaling another nucleus, then the two nuclei have identical morphology. Since our individual nuclei image masks were already centered and at the same scale, the only morphology-invariant transformation we disentangled with Harmony was orientation. Such disentanglement ensures that the modeled morphology is unaffected by orientation.

Given an input image of a nucleus, Harmony disentangles two key factors: semantic content and orientation. The semantic content captures the underlying information independent of how the nucleus is oriented, while the orientation component reflects its spatial rotation. During training, Harmony learns to represent and manipulate these factors simultaneously by adjusting the orientation while preserving the semantic content. To enforce consistency across different views, a dual-branch training strategy is employed, where both the original and a randomly rotated version of the same image are processed in parallel (Figure 1D). The system is trained to generate a consistent reconstruction, ensuring that the semantic representation remains stable regardless of orientation.

We obtained morphology quantification by clustering the semantic latent space learned by Harmony (Figure 1E). To ensure that the resulting semantic latent representations learned by Harmony properly represent the morphology information, we incorporated two significant modifications to Harmony training. The original Harmony [15] method uses a linear combination of two different loss functions: (1) reconstruction loss among the transformed and decoded images for both branches, and (2) KL divergence loss between the semantic content learned in both branches. We kept the reconstruction loss as it is. However, we replaced the KL divergence loss with a contrastive loss to ensure better morphological clustering. To calculate the contrastive loss, we used the Bhattacharyya coefficient instead of KL. Furthermore, we added another latent neighborhood loss component that ensures the semantic latent representations learned by Harmony actually correspond to the nuclear morphological variations. Details on these loss components can be found in Methods section.

After training the Harmony network with these two additional losses, the learned semantic latent factor *ϕ* contains high-fidelity morphology information. We obtained the semantic latent factor for all the nuclei images in the dataset and clustered them with a Gaussian Mixture Model (GMM). Thus, we obtained the nuclear morphology labels for each nucleus image. We performed several morphological image processing operations to obtain other nuclear attributes (nucleoli count, nuclear area, etc.) from the images (details in Methods).

To determine cell phenotypes, we used data from [18] that includes the grain counts of the RNAscope probe using the Halo software package by Indica Labs [19]. We aligned the grain counts with our segmentation results, achieving the grain count per nucleus for all the nuclei we segmented. As suggested in [18], in mice, if the grain count of TH per cell was at least 3, it was considered a dopaminergic cell (TH +) and if the grain count of VGLUT2 per cell was at least 1, it was considered a glutamatergic cell (VGLUT2 +). In humans, the threshold for TH and VGLUT2 was 5 and 3, respectively.

After determining the nuclear morphology labels for all nuclei and having associated information about the cell phenotype, age, and species, we performed statistical modeling to assess the effect of cell phenotype, age, and species on nuclear morphology. To this end, we statistically estimated whether any particular morphology label was increased or depleted in any particular cell type, age group, or species or in a combination of them. We estimated the proportion of morphology labels in each group based on cell phenotype or age. We assumed that the proportion outcome for each label follows a binomial distribution. Consequently, we fit a Bernoulli (logistic) mixed-effects model for each of the morphology classes. We then performed Type-II ANOVA test on each of these mixed-effects models and obtained p-values. On the basis of the p-values, we assessed the significance of the change in the morphology class of particular nuclei for any fixed effect.

### Nuclear morphology modeling and identification in a synthetic dataset

To validate our morphology modeling and identification pipeline, we first applied it to a synthetic dataset with known nuclear morphologies. Our synthetic dataset contains 20,000 images of nuclei with 20 different morphologies. The 20 different morphologies are sampled from [22] with varying eccentricity and irregular shape (Figure 2A). Among the 20 different morphologies, some are more similar to each other, whereas others are more dissimilar. Our task was to successfully model and identify the morphologies through the embedding space. We also tested embedding learning methods such as PCA, UMAP, t-SNE, autoencoder, and recently developed O2-VAE (orientation-invariant VAE) [22] to compare with our method in this task. For all the methods, we used a latent dimension of 2 for fair comparison and latent space interpretability.

**Figure 2:**
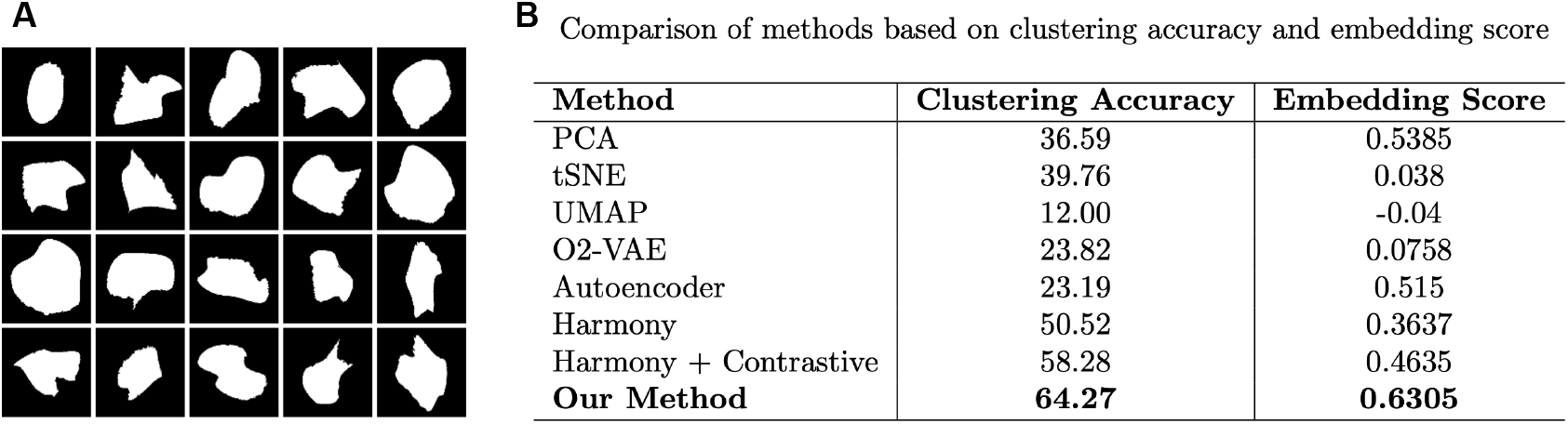
Our method largely outperforms other methods in morphology-identification in Synthetic dataset. **A**. The 20 different (based on eccentricity and randomness in shape) nuclear morphologies used in our dataset. **B**. The quantitative evaluation of our method and others, based on clustering accuracy and embedding score.

We quantitatively evaluated the performance of the baselines and our method on the simulated data with 20 different morphologies. We used two metrics for evaluation. First, we used the clustering accuracy in the latent space. We obtained the latent embeddings for all the images in our synthetic dataset using our method and baseline methods. We then performed clustering of the latent embeddings with a Gaussian Mixture Model (GMM) with 20 components. We then used Hungarian matching [23] to match the GMM clusters with ground truth classes and then computed the accuracy. We reported the accuracies obtained by the methods in Figure 2B in percentage. We also assessed the contribution of our contrastive loss and neighborhood loss, and found both of them to significantly contribute to the improvement in clustering accuracy.

We further evaluated the quality of the latent embeddings using a newly proposed metric called the ‘embedding score’. This metric quantifies the extent to which similarities in the image space align with similarities in the latent space, thereby assessing the morphology-awareness of the embeddings. Specifically, we computed this score by taking pairs of images along with their corresponding latent embeddings, calculating the structural similarity (SSIM) between the image pairs and the cosine similarity between the embedding pairs, and then estimating the correlation between these two similarity measures.

SSIM was used to measure image similarity and was implemented using the scikit-image package in Python. Both image similarity and embedding similarity values were normalized to fall within the range [0, 1]. To compute the correlation between them, we used the Pearson correlation coefficient from the scipy package in Python. The resulting correlation value was reported as the embedding score, which ranges from –1 to 1, where 1 indicates perfect positive correlation, –1 indicates perfect negative correlation, and 0 indicates no correlation.

Figure 2B shows that our method obtained the highest quality embeddings based on the embedding score. The incorporation of contrastive loss and neighborhood loss both significantly improved the embedding quality for nuclear morphology modeling.

### Cell type-specific nuclear morphology analyses across aging in mice

After successfully validating our method on synthetic data, we applied our pipeline to FISH data from mice (n = 15) with multiplex RNAscope expressions of the TH and VGLUT2 markers to perform cell-type and aging-specific nuclear morphology analyses in mice. For each mouse, we had one RNAscope data that imaged the whole-brain tissue (details in Supplementary Note 1). Our nuclei segmentation on the DAPI-stained channel of the data resulted in 1.1 million nuclei. Among them, 15,915 were in the manually marked midbrain region of interest. We trained our morphology identification method on the dataset. We performed Gaussian Mixture Model (GMM) clustering on the obtained morphology-aware latent space and created 10 non-overlapping clusters (Figure 3A). We classified the nuclei image set into 10 clusters, since we did not expect to observe more than 10 distinct nuclear morphology groups for our images. We calculated the distance of each nucleus from the centroids of the 10 clusters and assigned each nucleus to one cluster. We refer to these morphology classes from **M0** to **M9**. Each nucleus has a probability of being assigned to any of the morphology classes, and we obtained this probability from our GMM clustering model. For each morphology class, we visualized the 10 highest probability samples (after automatic alignment using our Harmony-based model) in Figure 3B.

**Figure 3:**
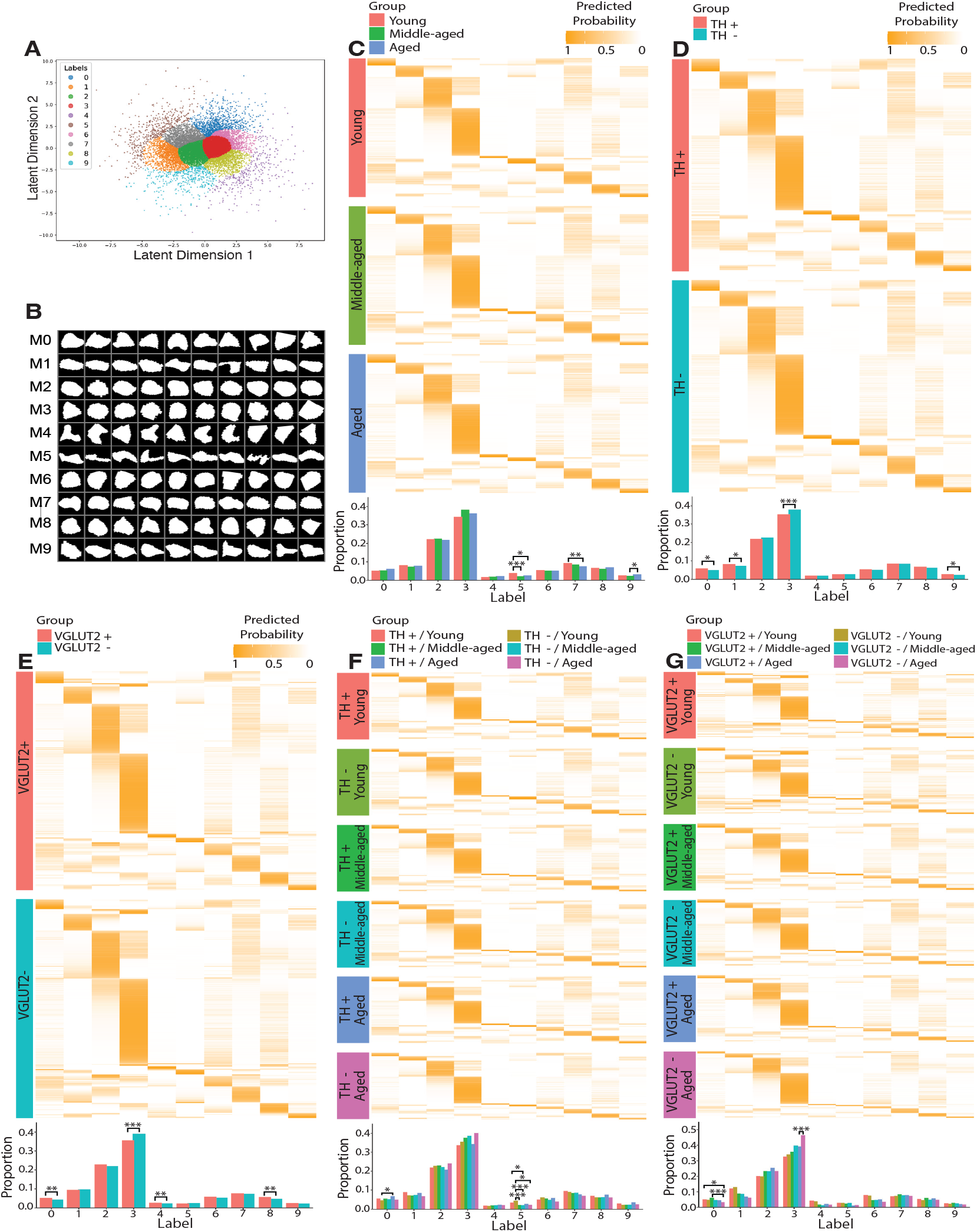
Nuclear morphology analyses results in mice midbrain. **A**. The morphology-aware semantic latent space obtained by our method. The latent space is clustered into 10 classes with GMM. **B**. 10 representative samples for each of the 10 morphology classes. The classes are numbered from *M* 0 to *M* 9.**C**. We plot the proportion of each nuclear morphology class (barplot in bottom) and the probability of each morphology class for all nuclei (heatmap in top) across three different age groups (young, middle-aged, and aged). **D**. and **E**. show similar plots across different cell phenotypes based on TH (**D**) and VGLUT2 (**E**), respectively. **F**. and **G**. show such a plot across different cell phenotypes and age groups, where **F**. and **G**. is based on TH and VGLUT2 respectively. In these plots, we mark (with *\**) the morphology classes that significantly differ in proportion across different groups. We determine significance based on p-values. If the p-value is less than 0.05 but above 0.01, we mark it with a single asterisk (*\**). If the p-value is less than 0.01 but above 0.001, we mark it with two asterisks (*\*\**). If the p-value is less than 0.001, we mark it with three asterisks (*\* * \**).

From the figure, we inferred semantic descriptions for each morphology cluster. **M2** and **M3** are both circular nuclei, being next to each other at the center of the latent space plot (Figure 3A). **M5** is the furthest nuclear morphology cluster with the most irregularly shaped nuclei. **M1, M9**, and **M7** are oval-shaped less-circular nuclei, where **M7** has the least irregular shape and **M9** has the most irregularity. **M0, M6**, and **M8** are all near-circular nuclei with some deviations. **M6** nuclei usually have one straight edge, **M8** nuclei have two straight edges, and **M0** nuclei has more than two straight edges. **M4** nuclei, which are few in number (Figure 3A), have an arrow-like nuclear morphology.

We tested whether any of these clusters were significantly increased or depleted in any specific age group, cell type, or their combination in mouse images. To this end, we conducted five statistical tests with respect to five independent variables: (1) age, (2) TH-based cell type, (3) VGLUT2-based cell type, (4) age x TH-based cell type, (5) age x VGLUT2-based cell type using the statistical approach mentioned in the overview.

We found that several nuclear morphology groups were significantly different in each statistical test (Figure 3F-G). We observed that the **M0, M1, M3**, and **M9** clusters were all significantly different between dopaminergic and non-dopaminergic neurons. Whereas **M0, M1**, and **M9** are significantly (*p-value < 0*.*05*) increased in dopaminergic neurons compared to nondopaminergic neurons, **M3** was very significantly (*p-value < 0*.*001*) depleted in dopaminergic neurons. Since M3 consists of completely circular nuclei, and **M0, M1**, and **M9** are all less circular or oval-shaped nuclei, it can be inferred that **dopaminergic neurons have more irregular shapes compared to nondopaminergic neurons in the midbrain of mice**. In the case of VGLUT2-based cell type, we found that M0, M4, and M8 morphology groups were significantly increased in glutamatergic neurons compared to non-glutamatergic neurons. On the other hand, the circular M3 morphology cluster was very significantly (*p-value < 0*.*001*) depleted in glutamatergic neurons compared to non-glutamatergic neurons. Since M0, M4, and M8 morphology groups contain nuclei with edges (M4 having an arrow-like shape), and M3 consists of fully circular nuclei, it can be said that **glutamatergic neurons have more edge-like irregular patterns in their shape compared to non-glutamatergic neurons in the mouse midbrain**.

While comparing across different age groups, we observed that elongated nuclei with tapered ends in **M5** are significantly enriched in younger mice neurons compared to middle-aged and aged mice. The younger mice also contain significantly more oval-shaped **M7** neurons than aged mice. These results suggest that **younger mice have more irregular nuclear morphology than aged mice**. This finding is aligned with the previous study [14] that indicated stiffer nuclear shapes in aged mice compared to younger ones through time-lapse image analyses.

When looking at the interplay of TH or VGLUT2-based neuron phenotype and aging, we found several interesting patterns. In the interplay of TH and healthy aging, we observed the enriched presence of elongated **M5** nuclei in younger mice compared to aged ones only for non-dopaminergic (TH-) neurons, not dopaminergic (TH+) neurons. Moreover, the nuclei cluster **M0**, which were significantly enriched in dopaminergic neurons compared to non-dopaminergic ones, were actually enriched in aged dopaminergic neurons than young non-dopaminergic neurons. On the other hand, somewhat opposite phenomenon was observed for VGLUT2. The nuclei cluster **M0**, which were significantly enriched in glutamatergic neurons (VGLUT2 +) compared to nonglutamatergic neurons (VGLUT2-), were actually significantly enriched in only young and middle-aged glutamatergic neurons compared to aged nonglutamatergic neurons. Moreover, the high statistical significance of enrichment of **M3** morphology in non-glutamatergic aged neurons compared to glutamatergic aged neurons, while the other being insignificant, suggests that **the presence of glutamate receptors affects aged mice more than younger mice**. All these findings highlight exciting novel insights that attract further in-depth investigation.

### Cell type-specific nuclear morphology analyses across aging in humans

To investigate nuclear morphology between cell phenotype and different age groups in humans, we applied our pipeline to human midbrain FISH data (n=12) with multiplex RNAscope expressions of the TH and VGLUT2 markers. Unlike mice, where we had a whole-brain image of a single subject, we had several midbrain regions assayed for each individual human subject. For 12 human subjects, we had 876 regions in total. Further details on the data can be found in Supplementary Note 1. Our segmentation of the DAPI-stained channel of the data resulted in 58,514 nuclei. We trained our morphology recognition pipeline on this dataset and obtained 10 nuclear morphology labels (Figure 4A). We refer to these morphology classes from **H0** to **H9**. Each nucleus has a probability of being assigned to any of the morphology classes, and we obtained this probability from our GMM clustering model. For each morphology class, we visualized the 10 highest probability samples (after automatic alignment using our Harmony model) in Figure 4B.

**Figure 4:**
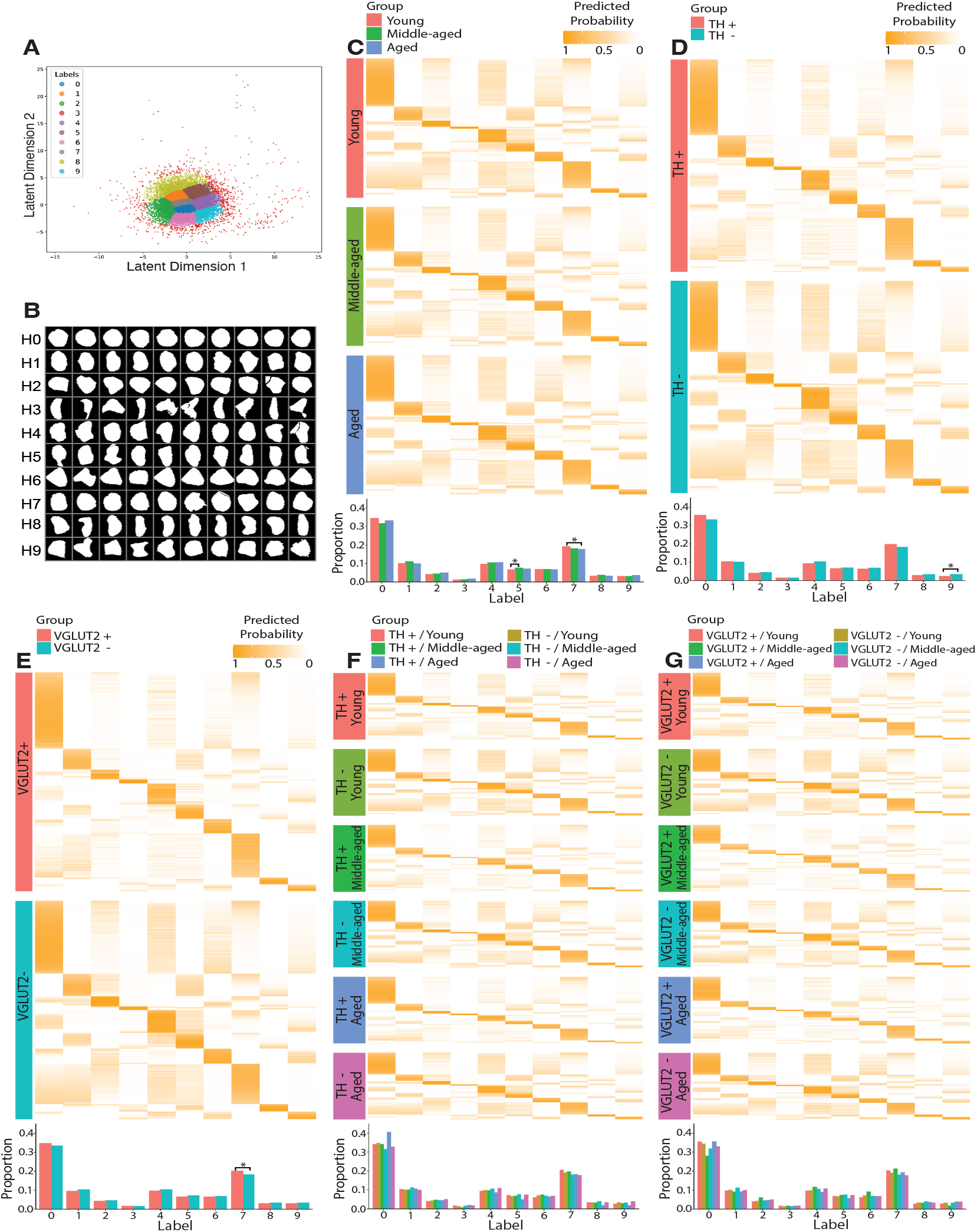
Nuclear morphology analyses results in human midbrain. **A**. The morphology-aware semantic latent space obtained by our method. The latent space is clustered into 10 classes with GMM. **B**. 10 representative samples for each of the 10 morphology classes. The classes are numbered from *H*0 to *H*9.**C**. We plot the proportion of each nuclear morphology class (barplot in bottom) and the probability of each morphology class for all nuclei (heatmap in top) across three different age groups (young, middle-aged, and aged). **D**. and **E**. show similar plots across different cell phenotypes based on TH (**D**) and VGLUT2 (**E**), respectively. **F**. and **G**. show such a plot across different cell phenotypes and age groups, where **F**. and **G**. is based on TH and VGLUT2 respectively. In these plots, we mark (with *\**) the morphology classes that significantly differ in proportion across different groups. We determine significance based on p-values. If the p-value is less than 0.05 but above 0.01, we mark it with a single asterisk (*\**). If p-value is less than 0.01 but above 0.001, we mark it with two asterisks (*\*\**). If p-value is less than 0.001, we mark it with three asterisks (*\* * \**).

It can be observed that **H0** morphology class, which is in the center of the latent space plot (Figure 4A), consists mainly of fully circular nuclei. In other words, **H0** is characterized by the highest roundedness. On the other hand, **H3**, which lies farthest from **H0** in the latent space (Figure 2a), has the most irregular shapes. **H7** and **H2** follows **H0** in terms of roundedness, where **H7** has subtle irregularities (one edge) and **H2** has more (two edges). **H2** and **H6** are very similar and also adjacent in the latent space. **H1, H4, H5**, and **H8** all contains oval-like nuclei with subtle differences between clusters. Among these four, **H1** and **H4** have higher roundedness than **H5** and **H8. H8** is the most oval-like shape, where **H1** is closest to circular **H2** and **H7**. The cluster **H6** and **H9** are similar and consist of somewhat circular nuclei with higher irregularities than **H2** and **H7**.

We tested whether particular morphology clusters were significantly increased or depleted in any specific age group, cell type, or their combination in human samples. To this end, we conducted five statistical tests with respect to five independent variables: (1) age, (2) TH-based cell type, (3) VGLUT2-based cell type, (4) age x TH-based cell type, (5) age x VGLUT2-based cell type using the statistical approach mentioned in the overview.

In general, we found fewer statistically significant patterns in humans than in mouse samples. However, at least one morphology cluster significantly differed between dopaminergic (TH+) and non-dopaminergic (TH-) neurons and between glutamatergic (VGLUT2+) and non-glutamatergic (VGLUT2-) neurons. For instance, **H9** cluster, which consists of circular nuclei with high irregularities, is significantly (*p-value < 0*.*05*) depleted in dopaminergic neurons compared to nondopaminergic neurons. On the other hand, **H7** cluster, which consists of circular nuclei with subtle irregularities, is significantly (*p-value < 0*.*05*) increased in glutamatergic neurons compared to non-glutamatergic neurons. However, the circular morphology was almost always increased in dopaminergic neurons compared to non-dopaminergic neurons and in glutamatergic neurons compared to non-glutamatergic neurons. In other words, **the presence of dopamine and the presence of glutamate provided a similar effect on nuclear morphology in human midbrain neurons**. This phenomenon was also observed in mouse neurons in our experiments with mouse data.

In terms of statistical significance with respect to age group, **H7** cluster, circular nuclei with subtle irregularities, gradually depleted from the young group to the middle-aged and from the middle-aged group to the aged group. The difference in the proportion of this cluster has been found to be statistically significant (*p-value < 0*.*05*) between the young and aged groups. Apart from **H7** cluster, **H5**, containing oval-shaped nuclei, is significantly (*p-value < 0*.*05*) increased in middle-aged group compared to young. However, it becomes depleted in the aged group again, suggesting no gradual aging pattern like **H7**. When examining age-specific cell populations within each phenotype (dopaminergic vs. non-dopaminergic, glutamatergic vs. non-glutamatergic), we did not observe statistically significant differences in specific morphology clusters (Figure 4F-G) as we did in mice. The different observations in mice and humans might not necessarily reflect species-specific effects on nuclear morphology, but could instead stem from discrepancies in data collection. In mice, the entire midbrain was imaged and all nuclei were segmented, providing comprehensive coverage. In contrast, only partial midbrain regions were imaged in human subjects, limiting the number of nuclei available for analysis. This restricted sampling may have reduced the statistical power, resulting in fewer significant findings. In future studies, imaging the full midbrain in human samples will help ensure more consistent and comprehensive comparisons across species.

We further investigated whether there are species-specific or brain-region specific nuclear morphology patterns across mice and human midbrain. However, we did not find any obvious significant patterns in such investigations (Supplementary Note 4). We also observed that several commonly measured nuclear attributes, such as nuclear area, nucleoli count, and average DAPI intensity, did not appear to notably vary with age or neuron phenotype in mice or humans (Supplementary Note 5).

## Discussion

Nuclear morphology has shown its potential to serve as a biomarker of neurodegenerative diseases and cellular senescence [1, 12, 7]. However, the relationship of nuclear morphology with neuron subpopulations with different phenotypes across aging has not been explored at a large scale due to a lack of effective automated pipelines to analyze such data. In this work, we investigated the effect of different neuron cell phenotypes (dopaminergic vs. nondopaminergic and glutamatergic vs. nonglutamatergic) on nuclear morphology and attributes in mice and human midbrains across different age groups from large-scale RNAscope image datasets. To this end, we segmented the nuclei from the DAPI channel of the RNAscope images in a semi-automated manner. We manually annotated several patches of the DAPI channel of the RNAscope image and fine-tuned the CellPose model [11] in the annotations. We used the fine-tuned model to infer segmentation masks for the DAPI-stained images. The segmentation results were further used to calculate nuclear morphology and other attributes (size, nucleoli count, etc.).

We identified an unmet opportunity to develop effective automated methods for modeling morphology from large-scale nuclei image datasets. Our analysis revealed that existing methods face challenges in distinguishing diverse nuclear morphologies when applied to datasets containing tens of thousands of nuclei with high morphological variability. To address this, we developed a novel method to model morphologies based on a transformation-disentangling auto-encoder network, called Harmony [15]. By leveraging the principle that morphology represents the intrinsic structure of objects independent of transformations, transformation-disentangling networks provide a compelling and intuitive framework for this task. Although recent work has explored transformation-invariant networks, these methods often lack interpretability over the transformation components, unlike transformation-disentangling networks. Moreover, we demonstrate that our method significantly outperforms a leading orientation-invariant autoencoder method [22], particularly in capturing the full spectrum of morphological diversity in synthetic datasets. We attribute this advantage to our model’s ability to bypass limitations associated with fast Fourier transform-based alignment steps used in the prior method [22].

The segmentation of nuclei in the RNAscope DAPI channel, a deep disentangled representation learning based nuclear morphology identification method, and downstream statistical analyses constitute our pipeline. While being highly effective for this study, the pipeline has several expected limitations in general. For example, our nuclear segmentation from the DAPI channel of the RNAscope images, though empirically had the best performance among other approaches, is still not perfect. Consequently, the segmentation may contain a few artifacts that are carried out through the downstream processes. However, with the availability of superior segmentation methods, artifacts from segmentation can be further reduced. Next, our deep learning-based nuclear morphology identification method may not be the most appropriate one to use for small-scale data. In fact, for small-scale datasets with a few hundred nuclear images with fewer morphology variations, traditional methods of pre-alignment and principal component analyses [24, 25] may provide sufficiently good results. In addition, our statistical analyses can also be further improved with bootstrap resampling or Bayesian mixed-effects models at the cost of substantial computational complexity.

Despite these limitations, our work is the first to systematically link nuclear morphology with cell phenotypes using RNAscope images, particularly in the context of aging in both mice and humans. Our computational approach is broadly applicable and can be extended to other RNAscope or FISH datasets to investigate how nuclear morphology relates to cell phenotypes across variables such as age, sex, disease states, etc. The segmentation model that we provide offers a practical solution for accurate nuclei segmentation in large DAPI-stained images. Furthermore, our method to learning and identifying nuclear morphology patterns can be easily adapted to study the morphology of other subcellular structures, such as mitochondria, the endoplasmic reticulum, and more, from fluorescence or electron microscopy data, without requiring much modification. Together, these contributions offer a flexible and scalable framework for integrating image-based transcriptomics with subcellular structural analysis.

Linking subcellular structural morphologies with distinct cell populations identified through RNAscope phenotypes offers important biological and clinical implications. One of the most compelling possibilities is the use of nuclear and subcellular morphologies as biomarkers for early disease detection. Changes in nuclear morphology within specific neuron subpopulations may signal the onset of neurodegenerative diseases such as Alzheimer’s disease, Huntington’s disease (HD), and Parkinson’s disease (PD), potentially enabling earlier diagnosis and intervention. Moreover, subcellular morphological features are closely tied to fundamental molecular processes, including epigenetic regulation and transcriptional activity. For example, nuclear morphology has been shown to correlate with the spatial organization of chromatin [26, 27]. By mapping nuclear morphologies to neuron subpopulations—defined by RNAscope phenotypes—it becomes possible to investigate population-specific chromatin-level alterations in the context of disease progression, cellular function, or response to therapeutic interventions. These insights open new avenues for understanding the relationship between cellular architecture and function, elucidating disease mechanisms at the level of specific cell types, and identifying new targets for drug development.

## Methods

### Nuclei Segmentation

We implemented a semi-automated pipeline to segment nuclei from the DAPI channel of the RNAscope images, addressing challenges posed by large image sizes and overlapping nuclei. The segmentation was performed leveraging existing segmentation models by finetuning them on several DAPI image patches that we manually annotated with Label-Studio [21].

#### Data Preprocessing for Segmentation

The initial dataset consisted of high-resolution images in TIFF format, which underwent several preprocessing steps to ensure consistency and enhance model performance. First, all images were converted to grayscale to emphasize intensity information, which is crucial for nuclei segmentation. Following grayscale conversion, pixel values were then normalized to a range of [0, 1] to facilitate stable convergence during model training. In addition, each TIFF image in the mouse dataset was divided into 64 patches, which were further subdivided into 16 subpatches to ensure effective manual annotation. All the subpatches were resized to 256 *×* 256 pixels to standardize the input dimensions across the entire dataset. For the human dataset, each TIFF image represents a patch and hence was directly used for segmentation. Then, several of the patches were manually annotated with LabelStudio [21] software.

#### Segmentation

We used our manual annotation to evaluate and fine-tune existing segmentation models. We split the annotated patches for training and testing sets. We used the UNet segmentation model in [28] and the CellPose model in [11]. We evaluated the performance of both models for nuclei segmentation in our test dataset. We used Intersection over Union (IoU) score as the evaluation metric. We found that both existing models demonstrated poor performance in our test dataset. Subsequently, we fine-tuned the U-Net and CellPose models on our manually annotated training dataset, which led to a significant improvement in the segmentation performance in the test set. Fine-tuning enabled the models to better adapt to our specific dataset, resulting in superior segmentation accuracy (Supplementary Note 2). We leveraged the fine-tuned CellPose model to infer segmentation masks for all DAPI images in our mouse and human datasets.

### Nuclear Morphology and Attributes Analysis

After nuclei segmentation, we proceeded with feature extraction and morphology analysis of the segmented nuclei. We also analyzed several nuclei attributes for the segmented nuclei, including nucleoli count, nuclei area, and average DAPI intensity. To further standardize the input for feature extraction, the segmented nuclei were processed as follows: using binary mask outputs by the segmentation model, the individual nuclei were cropped from the original images and resized to 3232 pixels. Since direct cropping often resulted in nuclei positioned close to the image boundaries, we uniformly added a 4-pixel padding around each resized image. This padding yielded final images of 4040 pixels, ensuring that the nuclei were placed centrally.

#### Nuclear Attributes Analysis

Following segmentation, we conducted a detailed analysis of the nuclei attributes, focusing on key metrics such as the number of nucleoli, nuclei area, and average DAPI intensity (which often is linked to transcriptional activity).

##### Nucleoli counting

Nucleoli counting was performed using a multistep image pre-processing and segmentation approach. Initially, background subtraction was carried out using a median filter with a kernel size of 7, effectively removing unwanted background elements and isolating the foreground signal. This was followed by signal normalization to a range of [0, 255], which improved the visibility of key features within the nucleus. To reduce noise and refine the signal, a Gaussian blur was applied.

For segmentation, we used color-based techniques by converting the images from the RGB color space to the HSV color space. This allowed for more precise segmentation based on color discrimination. The threshold values in the HSV space were chosen empirically, resulting in binary masks that isolated the nucleoli regions. Morphological operations, including opening and closing, were applied to clean the binary masks, ensuring that small noise elements were removed and discontinuous regions were joined. Finally, the number of nucleoli was determined by contour detection using the OpenCV findContours function, ensuring accurate counting.

##### Nuclear area and average DAPI intensity

The area of each nucleus was calculated by adding the number of pixels with a nonzero value within the binary mask. This provided a quantitative measure of nucleus size. The average DAPI intensity was calculated by summing the pixel intensities within the masked region in the original image.

### Nuclear morphology identification

We used a transformation-disentangled representation learning framework named Harmony [15] to identify nuclear morphology. Since our nuclei images were centered and at the same scale, we only disentangled orientation or planar rotation with our Harmony-based framework.

At a high level, Harmony [15] is an autoencoder-like framework. Taking an image sample *x* visualizing a single nucleus as input, Harmony [15] encodes a 2D rotation parameter *θ* and a semantic latent distribution *ϕ*, where *ϕ* contains the orientation-agnostic semantic information of the nucleus. The rotation parameter *θ* is used to rotate the nucleus image *x*. Simultaneously, *ϕ* is used to reconstruct the rotated nucleus image with a decoder. To ensure that the decoded images are in orientation-agnostic form, a siamese-like training is used where the same process is applied to a randomly rotated version of the nucleus image *x*. The network is trained in such a way that the decoder learns to reconstruct the same nucleus image for both branches (Figure 1D).

#### Model architecture and implementation

As an autoencoder-like architecture, Harmony consists of an encoder and decoder that are learned through training. The encoder consists of several (5 for mice and human data) fully connected layers with Leaky ReLU activations, which map the input image into a latent space. The final layer of the encoder produces the parameters for the semantic latent distribution. On the other side, the decoder network reconstructs an image from a latent vector sampled from the semantic distribution. The decoder mirrors the encoder in terms of architecture, using fully connected layers (5 for mice and human data) with Leaky ReLU activations. The final layer of the decoder applies a sigmoid activation function, which compresses the pixel values into the [0, 1] range.

The model was implemented in PyTorch. The parameters of the models were learned through gradient-based optimization. We trained the model with the AdamW optimizer with a minibatch size of 100 and a learning rate of 0.001. The model was trained for 100 epochs on a single NVIDIA A100 GPU. The loss function optimized during training is described below:

#### Loss function

Our model’s total loss function consists of three main components: the sum of squared error (SSE) reconstruction loss, the contrastive loss based on the Bhattacharyya coefficient, and the latent neighborhood loss.

#### Reconstruction loss

For the reconstruction loss in the autoencoder-like Harmony method [15], we use the sum of squared error (SSE) loss. Given that the model includes an autoencoder structure, the SSE loss measures the pixel-wise difference between the image resulting from rotating the input *x* by *θ* and the decoder output 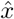 by the model:

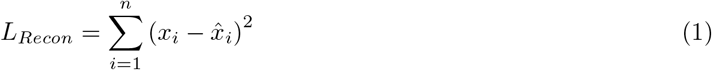

where *x*_*i*_ is a pixel in the rotated image and 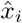 is the corresponding pixel in the decoded image, *n* is the number of pixels in the image *x*.

#### Contrastive loss with Bhattacharyya coefficient

The contrastive loss is designed to encourage the semantic latent representations of similar (positive) samples to be close together, while those of dissimilar (negative) samples are pushed apart. Here’s how positive and negative samples are defined and used in the model:

Positive Samples (**x**^+^): For a given single nucleus image *x*, the positive samples **x**^+^ are images that share certain similarities with *x*. In our approach, these positive samples are generated by applying random transformations to the input image *x*. These transformations ensure that the latent representation of the input and its positive samples are close to each other in the latent space, as they are essentially different forms of the same underlying image content. In this work, we used rotation as the transformation to generate positive samples.

Negative Samples (**x**^*−*^): Negative samples **x**^*−*^ are images that belong to a different class or content compared to the input image *x*. In our case, we performed batch training and the negative samples for the input image *x* are selected as the remaining images within the same batch **x**_**B**_. The goal is to ensure that the latent representations of these negative samples are pushed farther away from the representation of the input sample, reinforcing the separation between different types of data.

The contrastive loss is then defined as

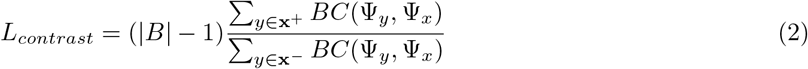

Here, *BC*(*ϕ*_*y*_, *ϕ*_*x*_) denotes the Bhattacharyya coefficient between the probability distributions of the latent representations *ϕ*_*y*_ and *ϕ*_*x*_, where *ϕ*_*x*_ and *ϕ*_*y*_ are semantic latent representations inferred by Harmony encoder for images *x* and *y* respectively. Details on the calculation of the Bhattacharyya coefficient are provided in Supplementary Note 3. |*B*| is the batch size during training. The loss encourages the latent representations of the positive samples to be close to the input while pushing the representations of the negative samples farther apart. Using random transformations for positive samples and selecting dissimilar images as negative samples, the model effectively learns to distinguish between similar and dissimilar contents in the semantic latent space.

To balance the difference in the number of positive and negative samples, we introduce the scaling factor |*B*|*−* 1, representing the proportion of negative to positive samples, ensuring stability in the training process.

#### Latent neighborhood loss

To preserve local neighborhood relationships of images in the input space in the semantic latent space, we incorporate a loss based on stochastic neighborhood embedding (SNE). This loss minimizes the divergence between pairwise similarities in both the high-dimensional image space and the low-dimensional latent space. The loss can be expressed as follows:

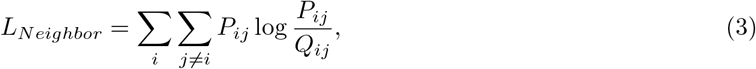

where *P*_*ij*_ is the probability that image *x*^(*i*)^ and *x*^(*j*)^ are neighbors in the high-dimensional image space, computed using a Gaussian kernel. *Q*_*ij*_ represents the probability that latent representations of *x*^(*i*)^ and *x*^(*j*)^ are neighbors in the low-dimensional latent space, computed using Student’s t-distribution (Supplementary Note 3). The loss function minimizes the difference between these distributions, thereby preserving neighborhood relationships and encouraging similar points in the high-dimensional image space to remain close in the latent space.

#### Total loss

The total loss for the model is a linear combination of the reconstruction loss, contrastive loss, and latent neighborhood loss.

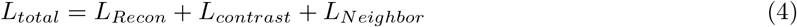

This combined loss ensures that the model produces latent representations that are semantically meaningful for morphology classification, well-structured for dimensionality reduction, and capable of accurate orientation-disentangled reconstruction.

#### Gaussian mixture model clustering in latent space

After extracting the two-dimensional semantic latent representations using Harmony, we applied a Gaussian Mixture Model (GMM) to cluster the latent space into *K* distinct classes. To apply GMM, we use the scikit-learn package in Python. The expectation maximization process in GMM was executed either for 100 iterations or until convergence with a convergence threshold of 0.01, whichever happened first. Each class represents a group of nuclei that share similar morphological characteristics based on their position in the latent space. This was qualitatively validated by visualizing the orientation-disentangled high-probability sample images within each class.

### Statistical analyses

We used the generalized mixed effects model (GLMM) to assess the effect of age, cell phenotype, or their combination on outcome variables such as nuclear morphology and other nuclear attributes. Since our data had a nested structure, we also considered random effects for different levels of nestedness. For the mouse data, there were two levels of nestedness: Subject *→* Samples. On the other hand, for human data, there were three levels of nestedness: Subject *→* Regions *→* Samples. Consequently, for statistical analyses on mouse data, we used the subject-level random effect. For human data, we used both subject-level and region-level random effects. The general mixed model for mouse data can be expressed as follows.

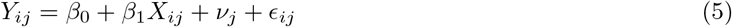

where *Y*_*ij*_ is the outcome variable for the i-th sample within j-th subject, *β*_1_ is the fixed effect for covariates (e.g., age, cell phenotype), *υ*_*j*_ is the normally distributed random effect for subject, and *epsilon*_*ij*_ is the residual error.

On the other hand, the general mixed model for human data can be expressed as follows:

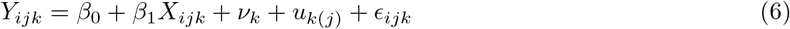

where *Y*_*ijk*_ is the outcome variable for the i-th sample within j-th region of the k-th subject, *β*_1_ again is the fixed effect for covariates (e.g., age, cell phenotype), *υ*_*k*_ and *µ*_*k*(*j*)_ are the normally distributed random effect for subject and regions nested within the subject, and *epsilon*_*ijk*_ is the residual error.

In these models, the outcome variable was either nuclear morphology or other nuclear attributes (nuclear area, nucleoli count, average DAPI intensity). For continuous-valued variables: nuclear area and average DAPI intensity, we considered Gaussian regression and applied a log transformation. For morphology classes, we used an indicator function to transform the ‘morphology label’ variable into a binary result. For the analysis of the label *l* where *Label* is the morphology label of a nucleus,

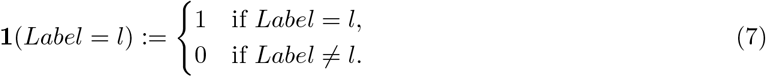

We assumed the outcome to be a binomial distribution as follows:

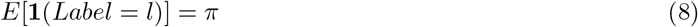

Where *Label* is the morphology label of *i*-th sample in the j-th subject (in mice) or *i*-th sample in the j-th region of the *k*-th subject (in humans).

We then fit a Bernoulli (logistic) mixed effects model taking logits of *π* as outcome variable *Y* : *Y* = *logit*(*π*).

For the count response (nucleoli count), we considered 3 types of regression models: Gaussian, Poisson, and negative binomial. While Gaussian regression is designed for continuous variables, Poisson and negative binomial regression are more suitable for counting data. After model fitting, we used the Akaike Information Criterion (AIC) to select the best model. A lower AIC value indicates a better model fit. Based on AIC values, we decided to use the Poisson model. We implemented the statistical models with lme4 package in R.

## Data Availability

The nuclear morphology quantification, and other nuclear attributes for the nuclei in the datasets are publicly available in https://doi.org/10.5281/zenodo.16752990.

## Code Availability

The code for morphology analyses and calculating nuclear attributes is publicly available in https://doi.org/10.5281/zenodo.16755197. This also includes the synthetic nuclear morphology dataset and a pretrained model.

## Acknowledgement

This work was supported by U.S. NIH grant R01GM134020.

## Author Contributions

M.R.U., Z.F., and M.X. conceived the research. M.R.U. designed and implemented the morphology identification method. M.R.U. designed the segmentation and nuclear attribute analysis method, and Z.Z., K.G., and Y.G. implemented these methods. R.C. and G.T. designed the statistical analyses. R.C. implemented the statistical analyses. J.K., S.B. J.G., Z.F. provided guidance on the mice and human datasets and feedback regarding the results on these datasets. M.R.U, Z.Z., and R.C. wrote the manuscript. All authors edited the manuscript.

## Competing interests

The authors declare no competing interests.

## Notes

### Competing Interest Statement

The authors have declared no competing interest.

