## Supplementary document for "Deep Disentangled Representation Learning Reveals Neuron Subtype-Specific Nuclear Morphologies Across Aging in Mice and Humans"

#### Supplementary Note 1: Mice and Human Dataset Description

**Mice Subjects:** Young (3-5 months old; N=5 males, 5 females), middle-aged (10-12 months old; N=3 males, 4 females), and aged (19-21 months old; N=4 males, 4 females) wild-type C57BL/6J mice (The Jackson Laboratory, Bar Harbor, ME; JAX#000664) were maintained on a 12:12 hour light:dark cycle with food and water available ad libitum. Age designations for the respective age-groups were determined according to previously described criteria28. Animals were euthanized with carbon dioxide followed by decapitation (for RNAscope fluorescent in situ hybridization experiments) or with isofluorane and transcardially perfused with 4% paraformaldehyde (for immunohistochemistry experiments). All experiments were approved by the University of Pittsburgh Institutional Animal Care and Use Committee. Animals were cared for in accordance with NIH animal care guidelines and ARRIVE guidelines for reporting animal research29-31. All efforts were made to ameliorate animal suffering.
**Human subjects:** Brain specimens (N=12 subjects) were obtained through the University of Pittsburgh Brain Tissue Donation Program following consent from next-of-kin, during autopsies conducted at the Allegheny County (Pittsburgh, PA) or the Davidson County (Nashville, TN) Office of the Medical Examiner. An independent committee of experienced research clinicians confirmed absence of lifetime psychiatric and neurologic diagnoses for all subjects based on medical and neuropathological records, toxicology reports, as well as structured diagnostic interviews conducted with family members of the deceased32, 33. Subjects were categorized as young (16-21 years, N=4), middle-aged (45-49 years, N=4), and aged (65-74 years, N=4). Each age group included two male and two female subjects, and subject groups did not differ in mean postmortem interval (PMI), RNA integrity number (RIN) or brain pH (Supplementary Table S1). One aged subject was removed from analysis due to outlier cell density levels of TH mRNA (TH+) cells and co-expressing TH and VGLUT2 mRNA (TH+/VGLUT2+) cells, as determined by the Robust Regression and Outlier (ROUT) method34, with a Q coefficient of 1 (Fig. S1). All analyses were therefore performed on the 11 remaining subjects, and the subject groups did not differ in mean PMI, RIN, or brain pH (all F2,8¡0.3 and all P¿0.75). All procedures were approved by the University of Pittsburgh's Committee for Oversight of Research and Clinical Training Involving Decedents and Institutional Review Board for Biomedical Research.
**Mice data:** We use RNAscope images from (N=15) mouse subjects where 5 were in the young group, 5 were in the middle-aged group, and 5 in the aged group. The images' dimensions are 19523 pixels (width) and 27890 pixels (height), capturing the whole brain section of mice. Each pixel is represented using 16 bits. Each image had four channels, each saved in a separate TIF file. There were separate sets of images for TH markers and VGLUT2 markers. The channel 0 of the images represent the DAPI-stained fluorescent image. The channel 3 represented the RNAscope probe for TH (in TH sets) or VGLUT2 (in VGLUT2 sets). There were no co-expression of TH and VGLUT2 markers.

All DAPI-stained images were used for nuclei segmentation, nuclear morphology identification, and attribute analyses. However, for TH and VGLUT2 markers, not all images had good signals. The images of only 12 mice had good TH or VGLUT2 signals - 4 from the young group, 4 from the middle group, and 2 from the aged group. Consequently, for neuron subtype-specific analyses in the midbrain, we only used images from these subjects.
**Human data:** We use RNAscope images from (N=12) human subjects, where 4 were in the young group, 4 were in the middle-aged, and 4 in the aged group. The image dimensions are 1024 pixels (width) and 1028 pixels (height), capturing several midbrain regions of the human subjects. There were 876 regions in total. Similar to mice images, each pixel is represented using 16 bits. There were separate images for DAPI stain, TH marker, and VGLUT2 marker. There were images for other housekeeping genes as well. All the images were in TIFF format. There were 3552 images in total. We used all the DAPI-stained images for nuclei

segmentation, morphology, and attribute analyses. All the TH and VGLUT2-stained images were used for neuron subtype-specific analyses.

#### Supplementary Note 2: Nuclei Segmentation Results

We perform nuclei segmentation on the DAPI-stained channel of the RNAscope images. To this end, we first used the Halo segmentation software by indica labs. However, this resulted in an undersegmentation of nuclei in the images S1A. Moreover, Halo is commercial software and can not be widely used by the community. To this end, we develop our own segmentation model.

We manually annotated several patches of the DAP-stained channel of the RNAscope images. We did our annotation using Label Studio. We first evaluated the performance of the existing segmentation models [1, 2] in our annotated data set. We found that these models, when applied directly for inference, do not provide sound segmentation results. However, once they are fine-tuned in our annotated training dataset, they perform significantly better in the test dataset. We perform 10-fold cross-validation so the chance of overfitting is less chance of overfitting and the superior performance means actual improvement in the segmentation task. We report the segmentation performance as the Intersection Over Union (IoU) score in Figure S1B. We can observe that the best segmentation performance in our test dataset is obtained by the fine-tuned CellPose [2] model, which we use as our final segmentation model to perform nuclei segmentation across all the RNAscope images in our dataset.

A

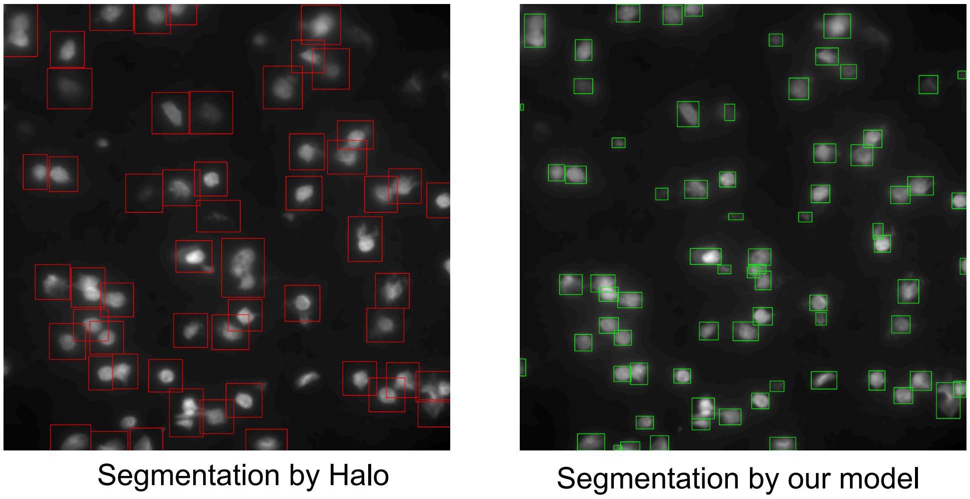


B

**Nuclei Segmentation Results**

| Models | IoU Score |
|---|---|
| Pretrained U-Net (Kromp et al.) | 0.49 |
| Pre-trained Cellpose (Stringer et al.) | 0.72 |
| Fine-tuned U-Net | 0.78 |
| Fine-tuned Cellpose | **0.84** |

Figure S1: **Nuclei Segmentation Results**. **A** A comparison between segmentation performed by Halo and our segmentation model. **B** Cross-validation segmentation results with different models.

Moreover, the IoU score is defined as:

$$\text{IoU} = \frac{\text{Area of overlap}}{\text{Area of union}}$$

**Where:**

- Area of Overlap = The area where the predicted and ground truth regions both exist in the nuclei segmentation mask.
- Area of Union = The total area covered by the nuclei segmentation masks (either predicted or ground truth, or both).

We also provided visualization of several segmentation results on the entire mouse brain. Figure S2.

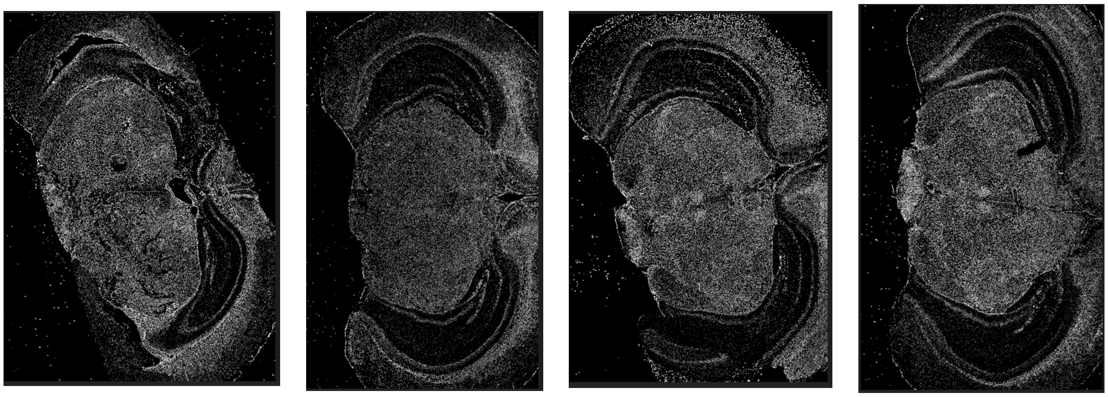

Figure S2: Nuclei segmentation results for several mice brain (whole brain section) DAPI-stained images.

### Supplementary Note 3: Loss functions in Nuclei Morphology Learning Method

#### Bhattacharyya distance

The Bhattacharyya distance is a measure of similarity between two probability distributions. It quantifies the overlap between the distributions, where a lower value indicates more similarity and a higher value indicates greater divergence.

For two probability distributions $P(x)$ and $Q(x)$ over the same domain $x$, the Bhattacharyya distance is given by:

$$D_B(P,Q) = -\ln\left(\sum_x \sqrt{P(x)Q(x)}\right) \tag{1}$$

where the **Bhattacharyya coefficient** is:

$$BC(P,Q) = \sum_x \sqrt{P(x)Q(x)} \tag{2}$$

Thus, the Bhattacharyya distance is:

$$D_B(P,Q) = -\ln(BC(P,Q)). \tag{3}$$

For continuous probability density functions (PDFs) $p(x)$ and $q(x)$, the Bhattacharyya coefficient is:

$$BC(P,Q) = \int \sqrt{p(x)q(x)}\,dx \tag{4}$$

and the Bhattacharyya distance is:

$$D_B(P,Q) = -\ln\left(\int \sqrt{p(x)q(x)}\,dx\right). \tag{5}$$

If $P$ and $Q$ are **univariate Gaussian distributions** with means $\mu_1, \mu_2$ and variances $\sigma_1^2, \sigma_2^2$, the Bhattacharyya distance simplifies to:

$$D_B(P,Q) = \frac{1}{4}\ln\left(\frac{1}{4}\left(\frac{\sigma_1^2}{\sigma_2^2} + \frac{\sigma_2^2}{\sigma_1^2} + 2\right)\right) + \frac{1}{4}\frac{(\mu_1 - \mu_2)^2}{\sigma_1^2 + \sigma_2^2}. \tag{6}$$

For **multivariate Gaussians**, the formula generalizes to:

$$D_B(P,Q) = \frac{1}{8}(\mu_1 - \mu_2)^T \Sigma^{-1}(\mu_1 - \mu_2) + \frac{1}{2}\ln\frac{|\Sigma|}{\sqrt{|\Sigma_1||\Sigma_2|}} \tag{7}$$

where $\Sigma = \frac{1}{2}(\Sigma_1 + \Sigma_2)$.

##### Neighborhood Embedding Loss

To make the semantic content inferred by Harmony correspond to the similarity and dissimilarities in the image space, we use a neighborhood embedding loss function. This loss function is based on t-SNE. In the following, we describe how we compute and implement this loss function.

**Pixel Space Similarities:** Given data points $x_i$ and $x_j$, we define a conditional probability $p_{j|i}$ that measures the similarity of data point $x_j$ to $x_i$:

$$p_{j|i} = \frac{\exp(-\|x_i - x_j\|^2/2\sigma_i^2)}{\sum_{k\neq i}\exp(-\|x_i - x_k\|^2/2\sigma_i^2)}$$

The bandwidth $\sigma_i$ is selected such that the perplexity of the conditional distribution equals a user-specified perplexity value. The symmetric joint probabilities are defined as:

$$p_{ij} = \frac{p_{j|i} + p_{i|j}}{2N}$$

**Embedding Space Similarities:** For the low-dimensional semantic embedding space counterparts $y_i$, the similarity $q_{ij}$ is computed using a Student's t-distribution with one degree of freedom (since low-dimensional):

$$q_{ij} = \frac{(1 + \|y_i - y_j\|^2)^{-1}}{\sum_{k\neq l}(1 + \|y_k - y_l\|^2)^{-1}}$$

**Optimization via KL Divergence:** We minimize the Kullback-Leibler divergence between the pixel-space and embedding-space distributions:

$$\mathrm{KL}(P\|Q) = \sum_{i\neq j} p_{ij}\log\left(\frac{p_{ij}}{q_{ij}}\right)$$

We optimize this loss using gradient descent.

#### Supplementary Note 4: Species-specific and brain region-specific nuclear morphology in mice and humans

We investigated whether there are species-specific nuclear morphology patterns across mice and human midbrain. To this end, we created a cross-species dataset that combined all nuclei masks obtained from mice and the human midbrain. From the latent space plot of morphology-aware semantic distributions obtained with our model, we observed no obvious morphological differences between human and mouse nuclei, suggesting the nuclear morphology is conserved in general across species for mouse and human samples.

In addition, we tested whether there are spatial region-specific nuclear morphology patterns for different cell phenotypes throughout aging in mice and humans. We plotted several heat maps showing the spatial distribution of nuclear morphology classes and cell phenotypes between different age groups in mice and humans (Figure S3A). For mice, we demonstrated the figures for the entire midbrain, and we did not observe any spatial region-based specificity for the morphology classes and the cell phenotype (Figure S3B). Since the entire midbrain was not imaged for humans, we plotted the morphology classes and cell types in the imaged region for humans (Figure S3C). Nevertheless, we did not observe any spatial patterns from the imaged regions. Both of these results suggest that nuclear morphology classes are not spatial region-specific and rather uniformly distributed across the midbrain or the midbrain region assayed in mice and humans.

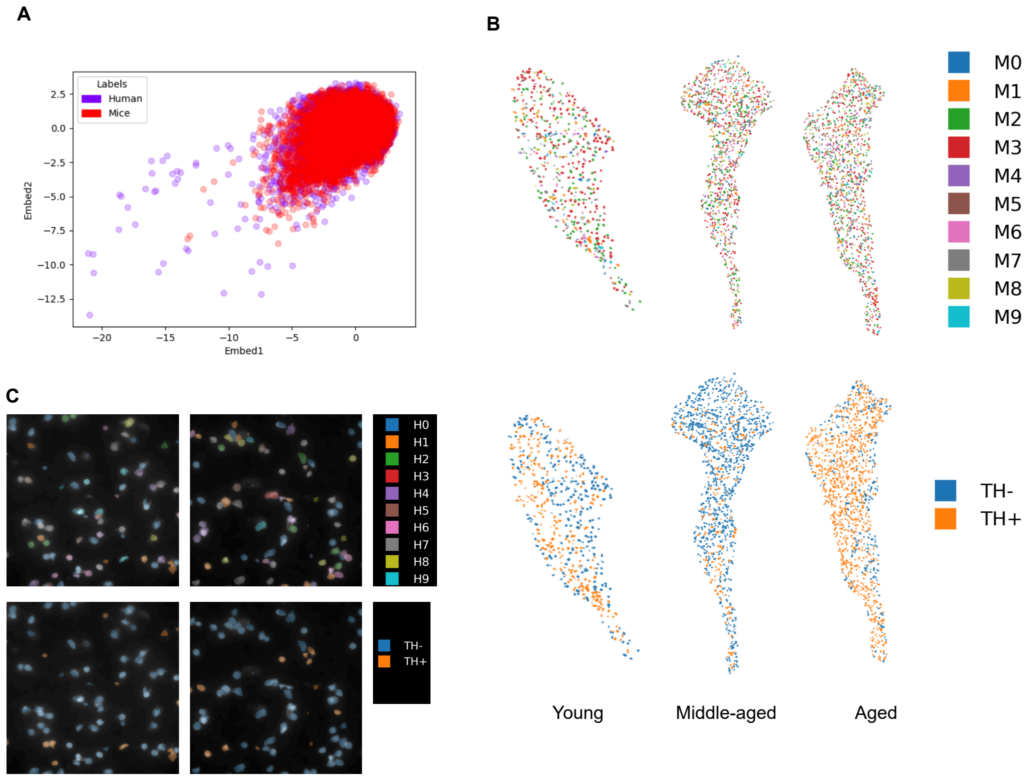


Figure S3: **A.** Morphology-aware semantic latent space visualization do not show differences between human and mouse midbrain morphologies. **B.** Visualization of mouse midbrain portions colored according to morphology labels (top) and TH-based cell phenotype (bottom). **C.** Visualization of two sample human midbrain assays colored according to morphology labels (top) and TH-based cell phenotype (bottom).

### Supplementary Note 5: Effect of Aging and Cell Phenotype on Nuclear Attributes

In addition to nuclear morphology, we also measured the effect of aging and cell phenotype on nuclear attributes such as area, nucleoli count, and average DAPI intensity (which has previously been linked to transcriptional activity of a cell). We plot the distributions of nuclear size, nucleoli count, and average DAPI intensity across different age groups and cell phenotypes based on TH (dopaminergic TH+ vs non-dopaminergic TH-) in mice midbrain in Figure S4(a). We plot the attributes in different age groups and cell phenotypes based on VGLUT2 (glutamatergic VGLUT2+ vs non-glutamatergic VGLUT2-) in mice midbrain in Figure S4(b). We provide similar plots for humans in Figure S5(a) and Figure S5(b), respectively.

From the plots, we do not see any obvious effect of aging or cell phenotype on nuclear area, nucleoli count, or average DAPI intensity. In addition, we performed several statistical analyses to measure the effect of aging and cell phenotype on these nuclear attributes.

Given the large sample size, we employed a generalized linear mixed model (GLMM) framework for statistical analysis. For nuclear area and average DAPI intensity—both continuous variables—we used Gaussian regression models. In contrast, nucleoli count is a discrete variable, for which a Gaussian model may be inappropriate. To identify the most suitable distribution for modeling nucleoli count, we compared Gaussian, Poisson, and Negative Binomial models based on their Akaike Information Criterion (AIC) scores. The model with the lowest AIC was selected as the best fit.

We provide the AIC scores in Table S1. In the mouse dataset, the Poisson model yielded the lowest AIC, while in the human data set, the negative binomial model performed best. Consequently, we used the Poisson model for mice and the Negative Binomial model for human data to assess the effects of aging and cell phenotype on nucleoli count.

For mouse data, we first used the nuclei $1,116,155$ throughout the entire mouse brain to assess whether aging has any effect on the nuclear attributes irrespective of cell phenotypes. We did not observe any

Table S1: AIC Scores for Nucleoli Count Distribution

| **Model** | **Mice** | **Human** |
|---|---|---|
| Gaussian | 3947245.37 | 304404.52 |
| Poisson | **3786828.37** | 312163.05 |
| Negative Binomial | 3786833.55 | **284854.11** |

statistically significant effect on the average DAPI intensity and nuclear area. For nucleoli count, we observed a statistically significant (p-value 0.01) reduction in younger group compared to middle-aged group. Though the reduction is significant, it is only by $\sim 6.6\%$. This suggests that although age may play a role in nucleoli count in whole mouse brain, it is not a major driver of variation.

After evaluating the effect of age across the entire mouse brain, we next focused on the mouse midbrain region to assess the influence of both age and cell phenotype. Cell phenotype annotations were available for only the midbrain region from 10 mice, comprising a total of 15,915 nuclei. When examining the impact of age on nuclear attributes within these midbrain nuclei, we found no statistically significant differences in nuclear area or average DAPI intensity. We observed only a marginally significant reduction in nucleoli count in young mice compared to middle-aged and old mice (p-value 0.0418), consistent with the trend observed in the whole brain analysis. However, the effect size was small, and the statistical significance was even weaker than that observed in the full-brain dataset. These findings suggest that age is not a major determinant of nucleoli count variation in the mouse midbrain.

Unlike age, which is a subject-level attribute, cell phenotype is defined at the sample (nucleus) level. As a result, while age remains constant across samples within a subject, cell phenotype can vary within the same subject. This within-subject variability contributes to a higher degree of freedom for cell phenotype compared to age. Possibly due to this higher degree of freedom, cell phenotype exhibits stronger statistical associations with nuclear attributes.

For example, we observed a statistically significant increase in nuclear area in dopaminergic (TH+) cells compared to non-dopaminergic (TH-) cells (p-value 0.0232). Similarly, dopaminergic cells showed a highly significant decrease in average DAPI intensity (p-value $< 10^{-16}$) and a modest but significant reduction in nucleoli count (p-value 0.0449).

Despite these statistically significant differences, the effect sizes were small, suggesting that cell phenotype may not be a major contributor to nuclear attribute variation in the mouse midbrain.

In human samples, we did not observe any statistically significant effect of age on average DAPI intensity or nuclear area. However, we detected a marginally significant increase in nucleoli count in young individuals compared to middle-aged and old individuals (p value 0.0497). Despite this subtle significance, the increase in nucleoli count was modest—approximately $\sim 14\%$—suggesting that age is not a major contributing factor behind the higher nucleoli count observed in younger human midbrain cells.

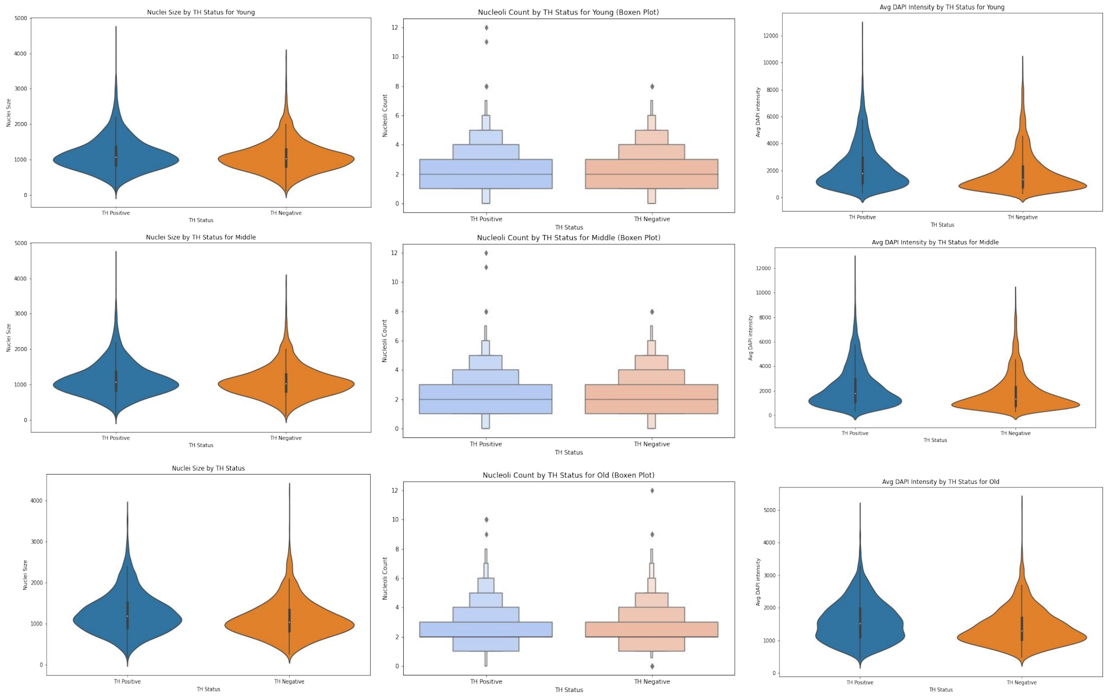


(a)

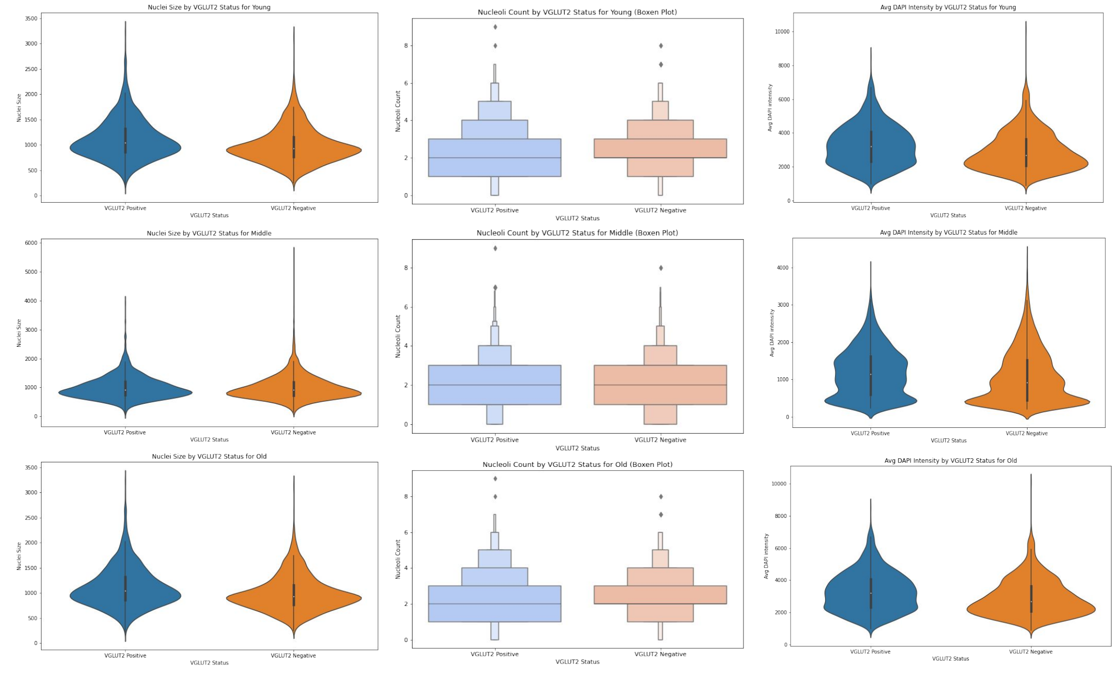


(b)

Figure S4: **a.** Comparison of nuclei size, nucleoli count, and average DAPI intensity in TH+ vs TH- nuclei in mice across different age groups. **b.** Comparison of nuclei size, nucleoli count, and average DAPI intensity in VGLUT2+ vs VGLUT2- nuclei in mice across different age groups.

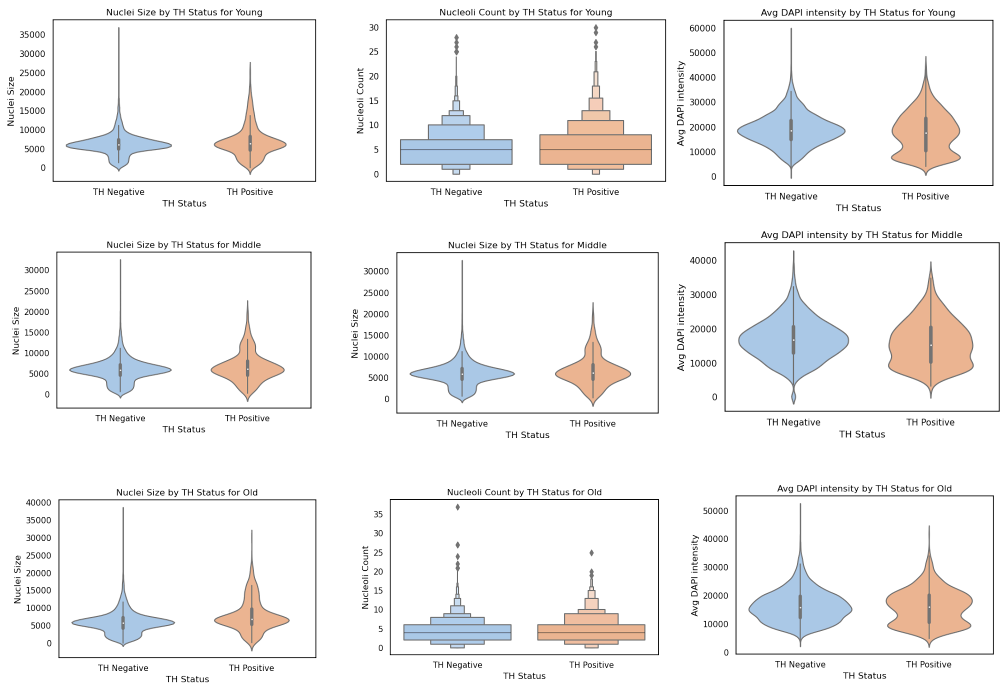


(a)

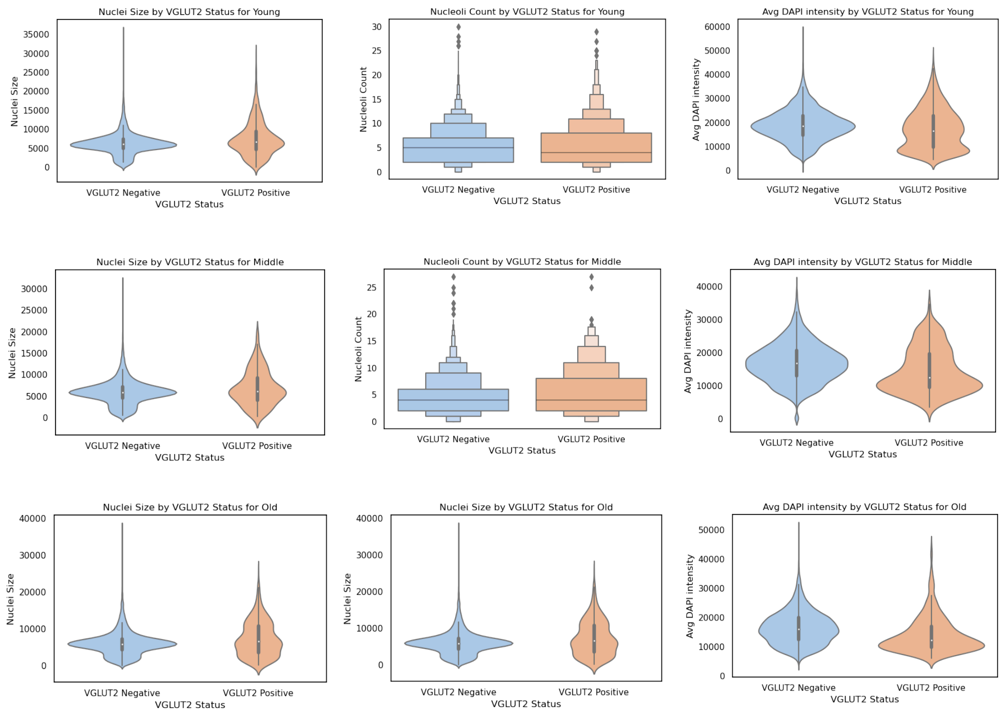


(b)

Figure S5: **a.** Comparison of nuclei size, nucleoli count, and average DAPI intensity in TH+ vs TH- nuclei in humans across different age groups. **b.** Comparison of nuclei size, nucleoli count, and average DAPI intensity in VGLUT2+ vs VGLUT2- nuclei in humans across different age groups.
